# Esco2 and Cohesin Regulate CRL4 Ubiquitin Ligase *ddb1* Expression and Thalidomide Teratogenicity

**DOI:** 10.1101/2020.09.02.280149

**Authors:** Annie C. Sanchez, Elise D. Thren, M. Kathryn Iovine, Robert V. Skibbens

## Abstract

Cornelia de Lange syndrome (CdLS) and Roberts syndrome (RBS) are severe developmental maladies that arise from mutation of cohesin (including SMC3, CdLS) and ESCO2 (RBS). Though ESCO2 activate cohesin, CdLS and RBS etiologies are currently considered non-synonymous and for which pharmacological treatments are unavailable. Here, we identify a unifying mechanism that integrates these genetic maladies to pharmacologically-induced teratogenicity via thalidomide. Our results reveal that Esco2 and cohesin co-regulate a component of CRL4 ubiquitin ligase through which thalidomide exerts teratogenic effects. These findings are the first to link RBS and CdLS to thalidomide teratogenicity and offers new insights into treatments.

## INTRODUCTION

Thalidomide, an over-the-counter drug used to relieve morning sickness during pregnancy in the late 1950s, leads to a suite of birth defects that include phocomelia, organ malformation, craniofacial abnormalities, and intellectual disabilities (*1*). These teratogenic effects result from inhibition of Cullin4 Ring Ligase (CRL4), the most common E3 ubiquitin ligase in eukaryotes and which contains Cullin4 (Cul4), DNA Damage Binding Protein 1 (Ddb1), and Ddb1-Cul4-Associated Factor (DCAF) Cereblon (Crbn) (*2, 3*). Inhibition of CRL4 function is solely responsible for thalidomide teratogenicity: development proceeds normally upon thalidomide exposure in embryos expressing thalidomide-resistant CRL4 and mutation of CRL4 subunits are sufficient to produce thalidomide-induced birth defects and intellectual disabilities (*4*). Roberts syndrome (RBS) and Cornelia de Lange syndrome (CdLS) are severe genetic maladies in which manifestations highly resemble those observed in thalidomide babies (*5, 6*). RBS arises through mutation of *ESCO2* while CdLS arises through mutation of cohesin subunits and regulators (*6*). ESCO2 acetylates SMC3 to activate cohesin, but CdLS and RBS etiologies are currently considered non-synonymous and for which pharmacological access is unavailable. Here, we document that SMC3 and ESCO2 knockdowns in zebrafish embryos provide robust models for CdLS and RBS and identify a unifying mechanism that integrates these genetic maladies to pharmacologically-induced teratogenicity via thalidomide. Our results reveal that Esco2 and cohesin co-regulate *ddb1* transcription, which is a component of CRL4 ubiquitin ligase through which thalidomide exerts teratogenic effects. These findings are the first to directly link both RBS and CdLS to thalidomide teratogenicity and transform current notions of cohesinopathies.

## RESULTS & DISCUSSION

### Conservation of phenotypes derived from *ESCO2* (RBS) and *SMC3* (CdLS) knockdowns, and Thalidomide exposure, in zebrafish embryos

Even though Esco2 acetyltransferase activates cohesin through acetylation of the cohesin subunit Smc3 (*7–9*), prevailing models of RBS and CdLS are quite different. Based on the initial discovery that Ctf7/Eco1 (herein Eco1, the homolog of human ESCO2) is critical for chromosome segregation (*10–12*), and that mutations in *ESCO2* lead to increase mitotic failure and apoptosis (*6, 13–16*), the prevailing model for RBS is based on mitotic failure that leads to proliferative stem cell loss (*16, 17*). Interestingly, CdLS cells typically do not exhibit increased mitotic failure or apoptosis, even though CdLS arises due to cohesin pathway gene mutations (*6, 18*). Based on the initial discovery that nipped-B (fly homolog of yeast Scc2 and human NIPBL - mutations in which produce the highest incidence of CdLS) plays a critical role in transcription regulation (*19, 20*), the prevailing model of CdLS is of transcription dysregulation (*6, 18*). Based on the phenotypic similarities between both RBS and CdLS genetic maladies, and the pharmacologically-induced birth defects resulting from in utero exposure to thalidomide, it became important to test whether all three are directly linked.

To address the role of Esco2 and cohesins in developing zebrafish embryos, we used *esco2* and *smc3’*-targetting morpholinos (MOs). Targeted protein knockdowns (KDs) of Esco2 and Smc3 were validated by Western blot of lysates obtained from MO-injected embryos. Lysates obtained from embryos injected with Standard control (SC) MO were used as control. A significant reduction of Esco2 and Smc3 protein levels in 24 hpf embryos were obtained with *esco2-ATG* MO and *smc3*-ATG MO respectively (Fig S1 A, B). These findings extend the previously validated target specificity and KD efficacy of both MOs in fin regeneration to embryonic development (*13, 21, 22*). Here, we assessed the role of Esco2 KD and Smc3 KD in developing zebrafish embryos. As a control, a standard control (SC) MO was used as it does not recognize target genes in zebrafish. MO injections were performed at the 1-cell stage and embryo phenotypes were assessed at 72 hours post fertilization (hpf). *esco2* MO injected embryos exhibited defects that include shorter body length (Fig. 1A, B), smaller eye size (Fig. 1A, C) and abnormal otolith development (Fig. 1A, D), compared to SC MO injected embryos. These phenotypes are consistent with the bone growth defects, small stature and hearing loss observed in RBS patients. *smc3* MO injected embryos, compared to SC MO injected embryos, also exhibited smaller body (Fig. 1A, B), reduced eye size (Fig. 1A, C) and a notable absence of otoliths within the otic vesicle (Fig. 1A, D), consistent with phenotypes present in CdLS patients. In combination, these results document that both Esco2 and cohesin are critical for proper craniofacial and body development in zebrafish embryos and provide robust models for RBS and CdLS phenotypes that arise during human development. In parallel, we exposed wild-type (WT) zebrafish embryos to 200μM, 400μM and 800μM concentrations of thalidomide, compared to DMSO treated control embryos (Fig. 2A). Significantly reduced body length was observed after treatment with 400μM and 800μM concentrations of thalidomide (Fig. 2A, B). Eye size was significantly reduced at all concentrations of thalidomide (Fig. 2A, C). Abnormalities in otolith development increased in a dose-dependent manner (Fig. 2A, D). These findings are consistent with thalidomide-dependent developmental defects (*1, 3*). The phenotypic overlap obtained from thalidomide treatment, and both Esco2 KD and Smc3 KD embryos, documents the efficacy of zebrafish RBS and CdLS models and spurred further efforts to ascertain the extent to which these pharmacological and genetic maladies are linked.

**Fig. 1.**
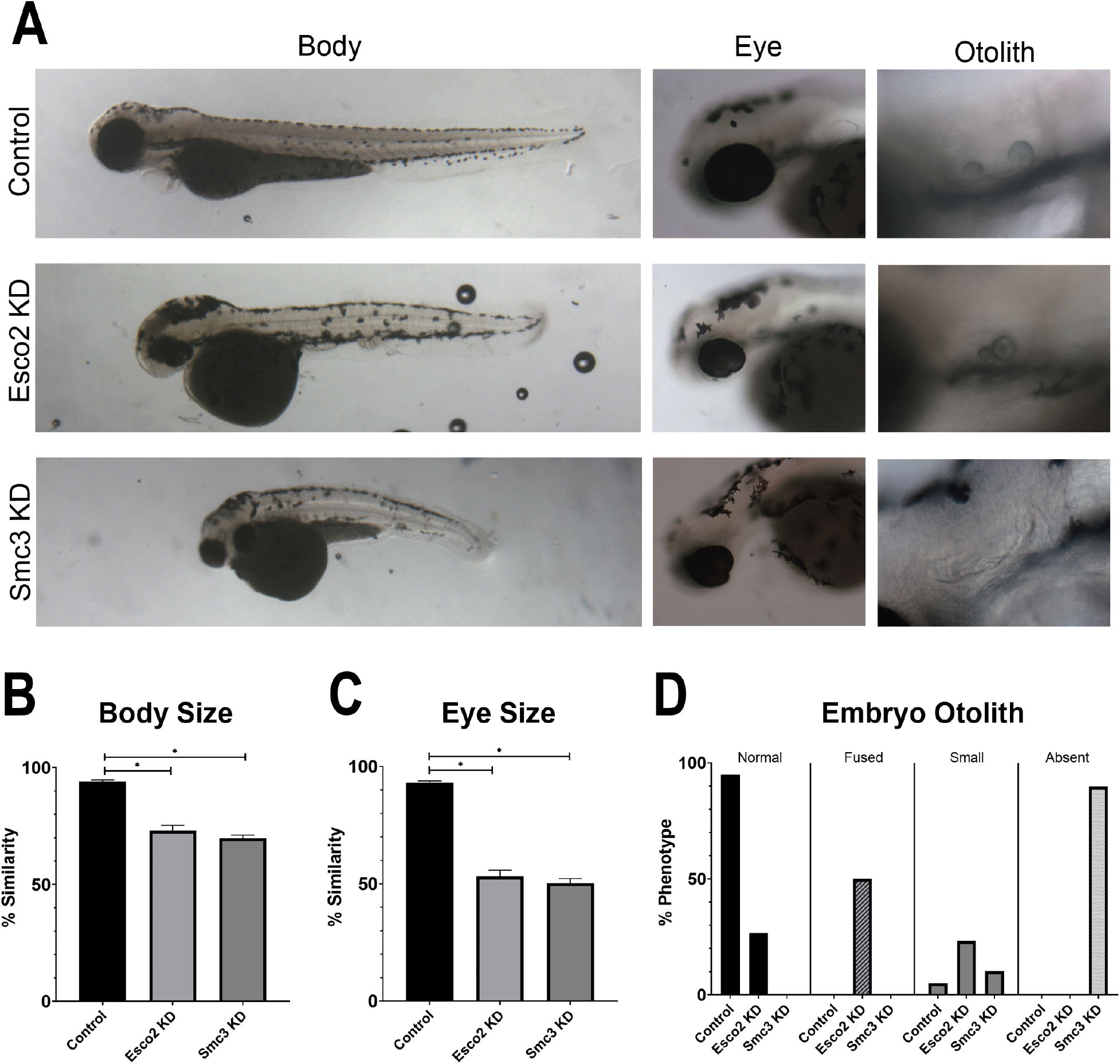
Esco2 KD and Smc3 KD phenotypes include reduced body and eye size, and an increase in abnormal otolith development. (A) Representative images of control embryos (SC MO injected), Esco2 KD (*esco2*-ATG MO injected) and Smc3 KD (*smc3*-ATG MO injected) embryos. For all experiments 24-40 replicates were analyzed and at least 3 independent trials were performed. (B) Quantification of body size from MO injected embryos were compared to un-injected WT embryos to obtain percent similarity. Graph reveals significant reductions of body length in Esco2 KD and Smc3 KD compared to control embryos (error bars represent s.e.m., one-way ANOVA with Turkey’s multiple comparison, *P<0.05). (C) Quantification of eye size from MO injected embryos were compared to un-injected WT embryos to obtain percent similarity. Graph reveals significant reductions of eye size in Esco2 KD and Smc3 KD compared to control embryos (error bars represent s.e.m., one-way ANOVA with Turkey’s multiple comparison, *P<0.05). (D) Graph shows percent of normal, fused, small, or absent otolith phenotypes with MO treatments. Data reveals 95% of control embryo otoliths exhibit normal phenotype, while Esco2 KD and Smc3 KD embryos exhibited 27% and 0% normal otolith phenotypes, respectively. An increase in abnormal otolith phenotypes was observed with KD treatments with predominantly fused phenotypes in Esco2 KDs and absent phenotypes in Smc3 KDs.

**Fig. 2.**
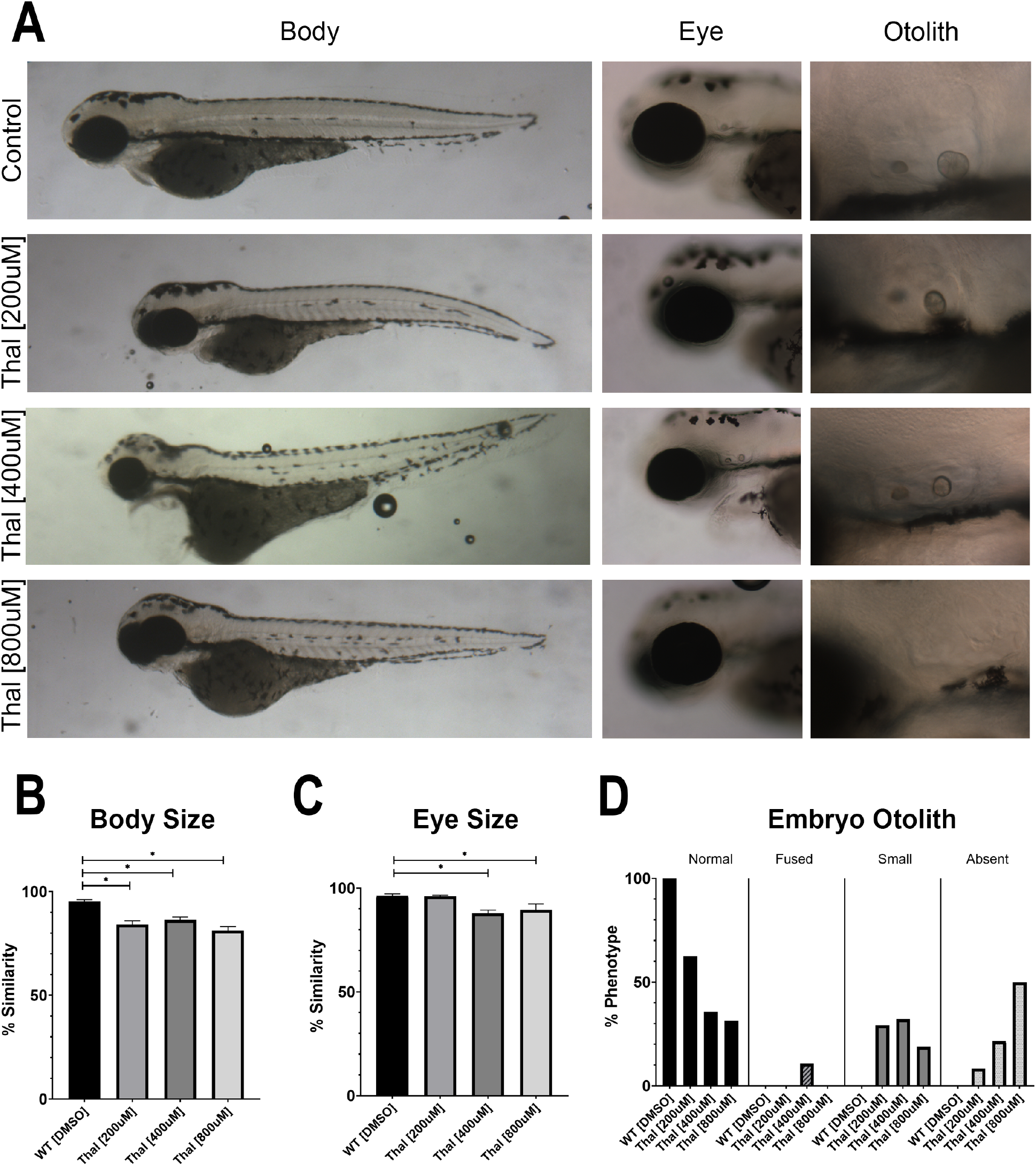
Phenotypes of thalidomide treated embryos overlap with Esco2 KD and Smc3 KD embryos. (A) Representative images of control embryos (WT treated with DMSO) and thalidomide (thal.) treatments (WT treated with 200μM, 400μM, and 800μM concentrations of thalidomide). For all experiments 16-28 replicates were analyzed and at least 3 independent trials were performed. (B) Quantification of body size after drug treatment were compared to untreated WT embryos to obtain percent similarity. Bar graph reveals a significant reduction of body length with thal. treatments compared to DMSO treated controls (error bars represent s.e.m., one-way ANOVA with Turkey’s multiple comparison, *P<0.05). (C) Quantification of eye size after drug treatment were compared to un-treated WT embryos to obtain percent similarity. Bar graph reveals a significant reduction of eye size with all thal. treatments compared to DMSO treated controls (error bars represent s.e.m., one-way ANOVA with Turkey’s multiple comparison, *P<0.05). (D) Graph shows percent of normal, fused, small, or absent otolith phenotypes with drug treatments. Data reveals 100% of control embryo otoliths exhibit normal phenotype, while 200μM, 400μM, and 800μM thal. treatments had 31%, 36% and 63% normal otolith phenotypes, respectively. An increase in absent otolith phenotypes was observed with increasing concentrations of thalidomide.

### Smc3 and Esco2 regulate expression of the CRL4 ligase gene *ddb1*

Thalidomide teratogenicity results from inhibition of CRL4 ubiquitin ligase function (*3*). Given the prevailing model that CdLS arises from transcription dysregulation (6, *18-20*), we tested for transcriptional deregulation of *cul4a, ddb1,* and *crbn* genes, each encoding a key component of CRL4 ligase, in Smc3 KD embryos. cDNA obtained from *smc3*-ATG MO injected embryos were assessed by qRT-PCR at 24 hpf and compared to SC cDNA. Fold changes in gene expression were calculated using Keratin as a housekeeping gene control. Neither *crbn* nor *cul4a* exhibited significant fold differences in gene expression. In contrast, *ddb1* was significantly reduced in Smc3 KD embryos (Fig. S2A). The prevailing model of RBS, based on mitotic failure and stem cell apoptosis (*l3-l7*), excludes a role for transcription dysregulation. Regardless, results document that *ddb1* expression is significantly downregulated in embryos injected with *esco2-ATG* MO at 24 hpf (Fig. S2A), similar to Smc3 KD embryos. These results provide compelling evidence for an emerging transcription dysregulation-based model of RBS (*6, 27, 28*) and link, for the first time, Esco2 and cohesin pathways to CRL4 regulation and thalidomide teratogenicity.

### Exogenous *ddb1* rescues severe growth defects associated with Smc3 Knockdown

If our finding that CRL4 is regulated by cohesin-dependent expression of *ddb1* is correct, consistent with the transcriptional model of CdLS etiology, then it should be possible to rescue Smc3 KD phenotypes by endogenous expression of *ddb1.* To test this hypothesis, embryos were injected with *smc3*-ATG MO, immediately followed by injection with *ddb1* mRNA (100ng/μL). Control embryos were solely injected with *ddb1* mRNA, which produced embryo development indistinguishable from un-injected WT embryos (Fig. S2B). When *smc3*-ATG MO injection was immediately followed by *ddb1* mRNA injection, however, developmental defects otherwise present in singly-injected *smc3*-ATG MO embryos were significantly reduced. For instance, both body and eye growth defects were significantly rescued by *ddb1* mRNA expression (Fig. 3A, B, C), and 70% of dual-injected embryos exhibited a normal otolith phenotype, compared to 0% normal phenotypes with *smc3* MO injection alone (Fig. 3A, D). Reduced levels (25ng/μL) of *ddb1* mRNA led to a partial rescue of Smc3 KD eye and otolith phenotypes, revealing a dosedependent response (Fig. S3A, B, C, D). These findings provide critical support for a model in which cohesins perform a transcriptional role upstream of CRL4 which, when abrogated, result in thalidomide-like teratogenicity.

**Fig. 3.**
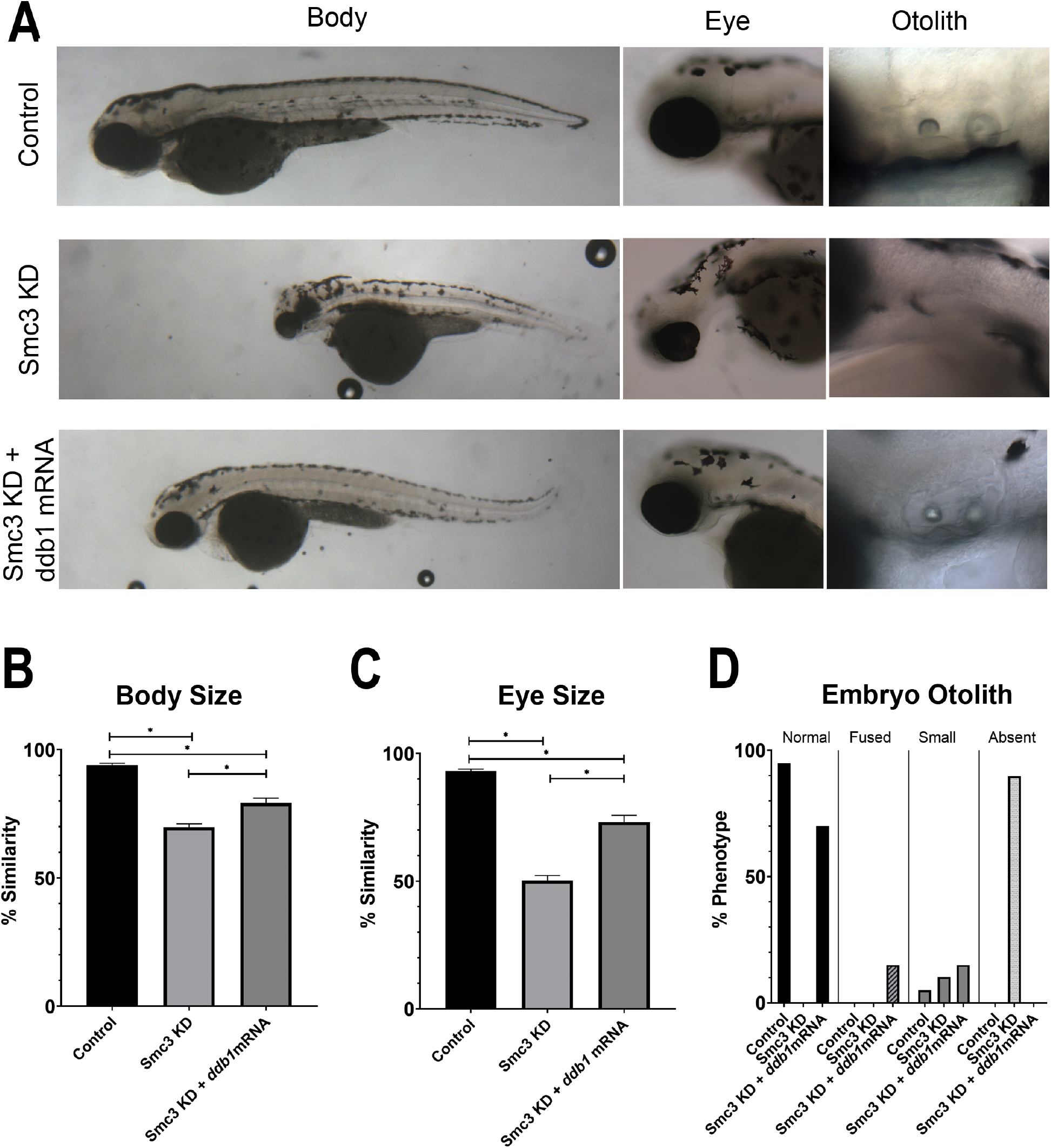
Exogenous *ddb1* overexpression rescues Smc3 KD phenotypes. (A) Representative images of control embryos (WT injected with *ddb1* mRNA), Smc3 KD (*smc3-* ATG MO injected) and Smc3 KD +*ddb1* mRNA (*smc3*-ATG MO co-injected with *ddb1* mRNA) embryos. For all experiments 26-40 replicates were analyzed and at least 3 independent trials were performed. (B) Quantification of body size from injected embryos were compared to un-injected WT embryos to obtain percent similarity. Bar graph reveals a significant rescue of body length in Smc3 KD +*ddb1* mRNA compared to Smc3 KD alone (error bars represent s.e.m., one-way ANOVA with Turkey’s multiple comparison, *P<0.05). (C) Quantification of eye size from injected embryos were compared to un-injected WT embryos to obtain percent similarity. Bar graph reveals a significant rescue of eye size in Smc3 KD +*ddb1* mRNA compared to Smc3 KD alone (error bars represent s.e.m., one-way ANOVA with Turkey’s multiple comparison, *P<0.05). (D) Graph shows percent of normal, fused, small, or absent otolith phenotypes with MO treatments. Data reveals 0% of Smc3 KD embryos exhibited normal otoliths, while 70% of Smc3 KD +*ddb1* mRNA embryo otoliths were rescued to normal levels. A decrease in absent otolith phenotypes was observed with *ddb1* mRNA co-injections compared to KD alone.

### Exacerbation of Esco2 knockdown phenotypes by exogenous *ddb1* reveals a feedback loop that is critical for development

We next tested for the effects of *ddb1* mRNA injection in *esco2*-ATG MO embryos. As opposed to rescuing *esco2*-ATG MO phenotypes, *ddb1* mRNA injection exacerbated the embryonic developmental defects. Body length and eye size were significantly decreased in the dual-injected embryos, compared to Esco2 KD alone embryos (Fig. 4A, B, C). Otoliths, if formed, were increasingly abnormal with a major fraction of embryos devoid of otoliths (Fig.4A, D). Reduced levels (25ng/μL) of *ddb1* mRNA co-injected with *esco2* MO, caused similar defects in body and otolith phenotypes compared to MO-only injected embryos, also revealing a dose-dependent response (Fig. S4A, B, C, D). Our findings are consistent with recent evidence that CRL4 targets and thus promotes Esco2 degradation (*23, 24*) and reveals a novel feedback loop through which Esco2 function may be regulated during development. Based on these findings, we favor a model in which *ddb1* mRNA injection counteracts Ddb1 reduction, caused by *esco2*-ATG MO, and elevates CRL4 activity. In turn, CRL4 upregulation decreases Esco2 levels and exacerbates Esco2 KD embryonic developmental defects.

**Fig. 4.**
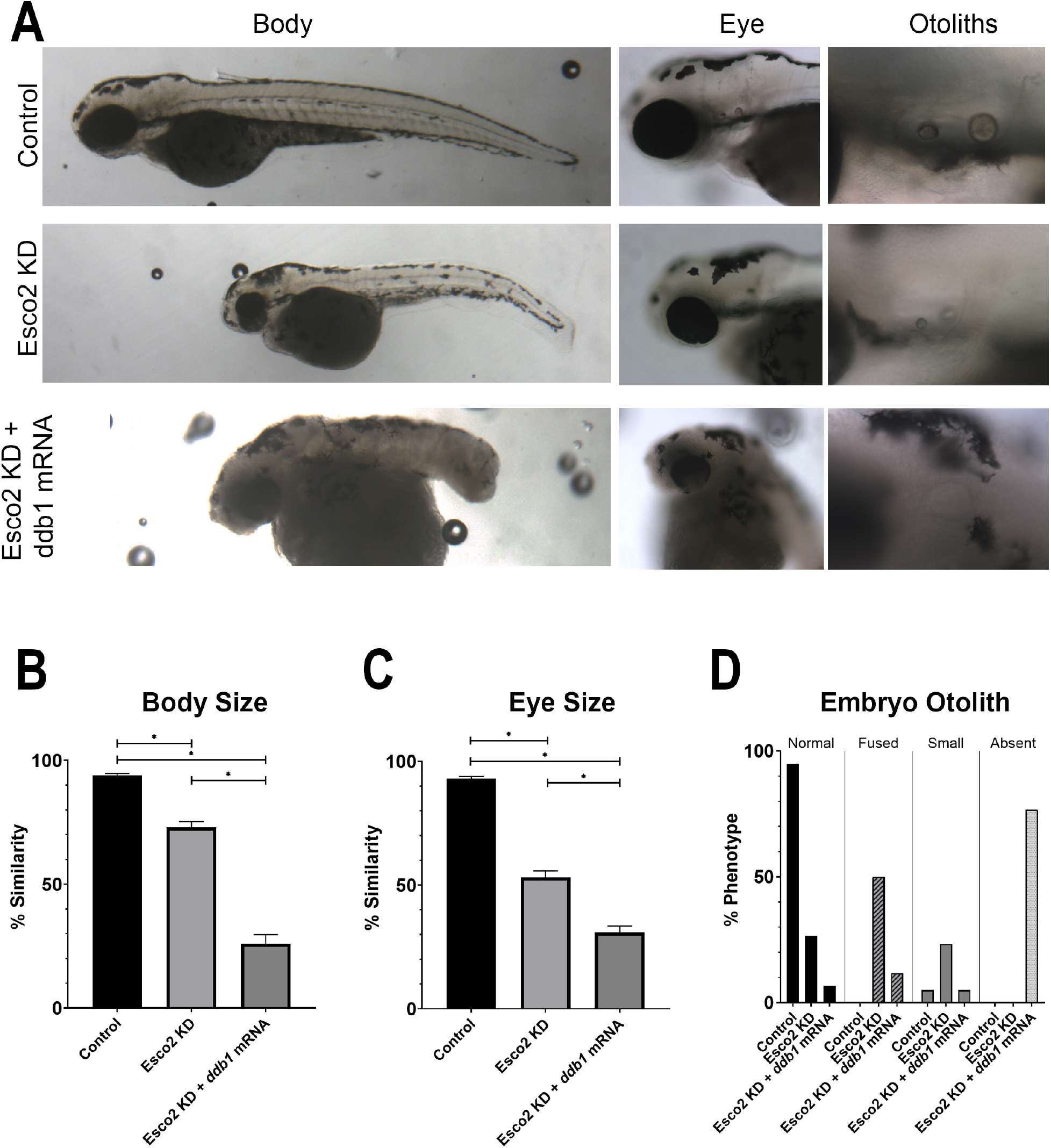
Exogenous *ddb1* overexpression exacerbates Esco2 KD phenotypes. (A) Representative images of control embryos (WT injected with *ddb1* mRNA), Esco2 KD (*esco2*-ATG MO injected) and Esco2 KD +*ddb1* mRNA (*esco2*-ATG MO co-injected with *ddb1* mRNA) embryos. For all experiments 29-38 replicates were analyzed and at least 3 independent trials were performed. (B) Quantification of body size from injected embryos were compared to un-injected WT embryos to obtain percent similarity. Bar graph reveals a significant reduction of body length in Esco2 KD +*ddb1* mRNA compared to Esco2 KD alone (error bars represent s.e.m., one-way ANOVA with Turkey’s multiple comparison, *P<0.05). (C) Quantification of eye size from injected embryos were compared to un-injected WT embryos to obtain percent similarity. Bar graph reveals a significant reduction of eye size in Esco2 KD +*ddb1* mRNA compared to Esco2 KD alone (error bars represent s.e.m., one-way ANOVA with Turkey’s multiple comparison, *P<0.05). (D) Graph shows percent of normal, fused, small, or absent otolith phenotypes with MO treatments. Data reveals 27% of Esco2 KD embryos exhibited normal otoliths, while only 7% of Esco2 KD +*ddb1* mRNA embryo otoliths were normal with an absent phenotype largely observed.

Discovering a pharmacologically-based mechanism through which genetic maladies, such as RBS and CdLS, converge represents a major advancement in our understanding of human development. Findings from this study suggest that the identification of CRL4 targets that act downstream of the Esco2-cohesin axis will profoundly impact both current models of birth defects and their treatment. In the former case, cohesins redistribute to over 18,000 new loci during zebrafish development (*25*), suggesting that regulating CRL4 subunit expression is only one example of many regulatory circuits through which Esco2 and cohesin function. A second example is *CX43,* a gap junction gene involved in skeletal development that is similarly under control of the Esco2-cohesin axis (*21, 22*). Mutations in *CX43* cause oculodentodigital dysplasia (ODDD) in humans and defects in bone segment regrowth in zebrafish (*6, 26*). The extent to which the Esco2-cohesin axis regulate genes independent of one another remains an important issue in development. In the latter case, the sea change in transforming RBS from one of mitotic failure to that of transcriptional dysregulation has profound implications for treatment. For instance, *ddb1* and *crbn* mutations result in elevated BK_Ca_ high conductance channel trafficking to the plasma membrane, increased ion conductivity, high neuronal excitation, and seizures (*27–29*). Paxilline, a BK_Ca_ channel blocker, reduces the incidence and severity of seizures (*28*). CLR4 knock-out mice also exhibit learning and memory deficits, an effect attributed to reduced translation of hippocampal glutamatergic synapse proteins via AMPK hyperphosphorylation. Compound C (an AMPK inhibitor) treatment normalizes glutamatergic protein levels and rescues both learning and memory deficits in *Crbn* knockout mice (*30*). The novel link between CRL4/thalidomide and RBS and CdLS is likely to extend to numerous developmental disorders, as well as cancers that are tightly correlated with cohesin mutations (*31, 32*). We look forward to future experiments that test the extent to which the molecular mechanisms revealed here provide for new strategies of treatment for a broad range of developmental maladies and cancers.

## MATERIALS AND METHODS

This study was performed in accordance with the recommendations in the Guide for the Care and Use of Laboratory Animals of the National Institutes of Health. These protocols were approved by Lehigh’s Institutional Animal Care and Use Committee (IACUC) (Protocol 187). Lehigh University’s Animal Welfare Assurance Number is A-3877-01.

### Morpholino (MO) Injections

MO purchased from GeneTools, LLC (Philomath, OR) were dissolved in sterile dH2O, for a 1mM concentration (sequences available upon request). These were heated to 65°C for 15 minutes prior to use. Full MO concentration resulted in embryo lethality, thus *smc3*-MO was diluted in 1X phenol red to a concentration of 0.5mM to allow for embryo comparisons at 72 hours post fertilization (hpf), and *esco2*-MO was diluted in 1X phenol red to a concentration of 0.25mM. A standard control (SC) MO with no target sequence in zebrafish was used as control. Microinjections were performed at the 1-cell stage using the Narishige IM 300 Microinjector and Nikon SMZ 800 for visualization. Zygotes were sorted for viability and fertilized embryos were kept in egg water and Ampicillin solution at 28°C. Embryos were de-chorionated using pronase if needed, then harvested for lysate or cDNA preparations or fixed in 4% paraformaldehyde (PFA), and kept at 4°C overnight for phenotype analysis. Embryos were stored in 100% methanol at −20°C for long term use after fixing.

### Thalidomide Treatments

As adapted from Ito et al. 2010 (*3*), a stock 400mM solution of thalidomide dissolved in DMSO was made in order to keep the final DMSO concentration under 0.1%. The 400 mM stock solution was diluted in E3 medium prewarmed at 65°C and mixed for 1 or 2 min to make 200μM, 400μM and 800μM final concentrations. Zebrafish embryos were manually dechorionated prior to thalidomide treatment by use of forceps. After chorion removal, embryos were immediately transferred to E3 medium containing thalidomide or DMSO only control and further incubated at 28°C until the 72 hpf timepoint was reached. E3 medium was replaced with freshly prepared medium every 12 hours. Embryos were fixed in 4% PFA and kept at 4°C overnight for phenotypes analysis. Embryos were stored in 100% methanol at −20°C for long term use after fixing.

### Embryo Lysates and Immunoblotting

MO injected embryo lysates were made at 24 hpf for Esco2, Smc3 and SC. Protocol was adapted from Schabel et al., 2019 (*33*). In short, embryos were de-chorinated with pronase then washed in E3 egg water. Individual embryos were placed in 1.5mL centrifuge tubes and all excess egg water was removed. 500μL of heptane were added then immediatly after 500μL of cold methanol were added and sample was fixed for 5 mins. Embryo was washed two times with 500μL of cold methanol then 2 times with 100μL of Embryo Buffer (EB). Embryos were homogenized in 20μL of EB. Three single embryo lysate preps were pulled to create one biological replicate. Lysates were stored at −80°C, 5X SDS Loading buffer was added and samples were boiled before use. A primary antibody specifically for zebrafish was used to detect Esco2 (1:1000, GenScript). Alexa 546 anti-rabbit (1:1000, Invitrogen) was used to detect Esco2 primary antibody. A primary antibody specifically for zebrafish was used to detect Smc3 (1:1000, Santa Cruz Biotechnology, sc-8198). Alexa 568 anti-goat (1:1000, Invitrogen) was used to detect Smc3 primary antibody. Mouse anti-α-tubulin (1:1000, Sigma-Aldrich, T9026) was used as a loading control. Alexa 647 anti-mouse (1:1000, Invitrogen) was used to detect the tubulin primary antibody. For measurement of band intensities, ImageJ software (https://imagej.nih.gov/ij/) was used. Relative pixel densities of gel bands were measured using the gel analysis tool in ImageJ software as previously described in Bhadra and Iovine, 2015 (*35*). Tubulin was used as a loading control and thus the relative expression calculations were based on the ratio of Esco2 or Smc3 to Tubulin.

### mRNA Rescue

Full length mRNA encoding for *ddb1* was designed using the sequence from the ZFin database. The plasmid was stored at −20°C until used. Plasmid was diluted to a 1:10 concentration of 0.2μg/μL in sterile water. One Shot Max Efficiency DH5αcells were transformed with the plasmid using standard procedures. The Qiagen Mini-Prep kit was used to isolate plasmid DNA from thetransformed bacteria. The plasmid DNA was then linearized by performing an AvrII digest. A transcription reaction was then performed using the Invitrogen mMessage mMachine kit. The concentration of the resulting mRNA was assessed using the Thermo Scientific Nanodrop 2000. This was also run on a formaldehyde gel and imaged using the BioRad Gel Doc. The mRNA was then diluted to concentrations of 25ng/μL and 100ng/μL in phenol red. Diluted mRNA was heated at 65°C for 5 minutes prior to injections into zebrafish embryos at the 1-cell stage as previously described. The mRNA was also co-injected into embryos that had been injected with the *smc3* and *esco2*-ATG start site blocker MOs. Both the mRNA injected embryos and coinjected rescue embryos were fixed at 72 hpf in 4% PFA overnight at 4°C and for phenotypes analysis. Embryos were stored in 100% methanol at −20°C for long term use after fixing.

### qRT-PCR

For qRT-PCR, total mRNA was extracted from around 15 embryos to make one biological replicate using Trizol reagent and the standard protocol. The resulting mRNA pellet was resuspended in a solution of DEPC H2O and RNAse Inhibitor, then the concentration of RNA was recorded using the Thermo Scientific Nanodrop 2000. For making cDNA, 1 μg of total RNA was reverse transcribed with SuperScript III reverse transcriptase (Invitrogen) using oligo (dT) primers. The resulting cDNA was diluted 1:10 for qRT-PCR, using the Rotor-Gene 6000. cDNA was made on *esco2*-ATG MO injected embryos, *smc3*-ATG MO injected embryos and SC MO injected embryos. For each cDNA used, primers at a 10μM concentrations were used. Three primers were specific for components of the CRL4 E3 Ligase affected by thalidomide and are available upon request. *keratin4* primers were used as a housekeeping gene control. For each PCR tube, 7.5μL of Sybr Green, 3μL 10μM Primers, 3.5μL sterile H2O, and 1μL cDNA was added. Analyses of the samples were done using Rotor-Gene 6000 series software (Corbette Research) and the average cycle number (C_T_) determined for each amplicon. Delta C_T_ (ΔC_T_) between housekeeping gene and CRL4 genes were calculated to represent expression levels normalized to *keratin* values. ΔΔC_T_ values were calculated to represent the relative level of gene expression and the fold difference was determined using the ΔΔC_T_ method (2^-ΔΔC^_T_) as previously described (*21, 34*).

### Imaging Analysis of Embryos

Zebrafish embryos fixed at 72 hpf were mounted on double cavity slides using 3% methyl cellulose for embedding. Embryo phenotypes were observed using the Nikon SMZ 1500, 1X objective at room temperature and the Nikon Eclipse 80i Microscope, 10X and 20X objectives at room temperature. Microscopes were equipped with SPOT-RTKE digital camera (Diagnostic Instruments) and SPOT software (Diagnostic Instruments) for image acquisition. The images obtained were then used to quantify whole embryo length, eye diameter, and otolith phenotypes. Percent similarities were obtained for body and eye measurements for each treatment compared to wild-type (WT) un-treated embryos. First, percent difference was calculated by taking the change in value between treated/injected embryo and un-treated/un-injected WT embryos, divided by the average of the numbers, all multiplied by 100. Percent similarities were then obtained by subtracting percent difference from 100. Otolith phenotypes were observed and scored into four categories: normal otoliths, fused otoliths, small otoliths, and absent otoliths. Otolith diameters were measured to distinguish between normal and small otoliths. Anterior and posterior otolith diameters were added; a sum less than 50μm was classified as small. Percent of embryos with each otolith phenotype were calculated by taking number of embryos in each category divided by total embryos analyzed.

### Statistical Analysis

ANOVA tests were used to determine if there was a statistically significant difference in body size, and eye size between knockdowns (KD) and controls. Statistical analysis was performed using ordinary one-way ANOVA tests. Two-tiled paired t-tests were used to determine if there was a statistically significant difference in qRT-PCR analysis between KDs and controls. N values of at least 16 were used in every experiment. Only values giving P < 0.05 are reported.

## SUPPLEMENTAL FIGURES

**Fig. S1.**
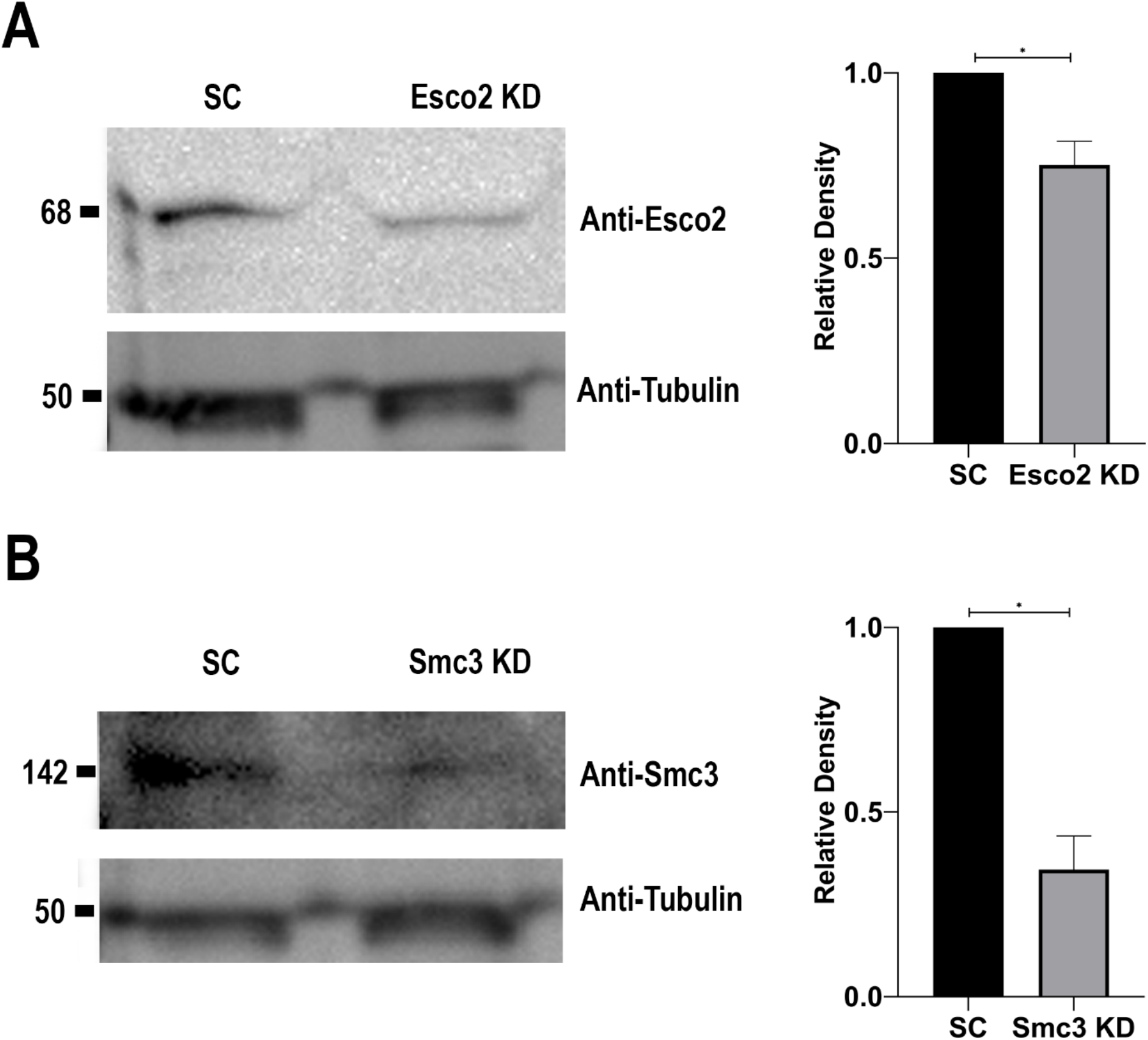
Validation of Esco2 and Smc3 protein reduction with MO injections (A) Western blot of SC embryo lysates compared to *esco2* MO embryo lysates. Immunoblots were probed with Esco2 and Tubulin antibodies. Analysis detects Esco2 at a predicted size of 68 kDa and Tubulin at the predicted size of 50 kDa. Graph shows the average relative densities of the Esco2 bands between experimental and control sample from 3 biological replicates. Esco2 protein levels are significantly reduced by 25% with MO injection compared to SC injected embryos (error bars represent s.e.m., un-paired t-test, *P<0.05). (B) Western blot of SC embryo lysates compared to *smc3* MO embryo lysates. Immunoblots were probed with Smc3 and Tubulin antibodies. Analysis detects Smc3 at a predicted size of 142 kDa and Tubulin at the predicted size of 50 kDa. Graph shows the average relative densities of the Smc3 bands between experimental and control sample from three biological replicates. Smc3 protein levels are significantly reduced by 66% with MO injection compared to SC injected embryos (error bars represent s.e.m., un-paired t-test, *P<0.05).

**Fig. S2.**
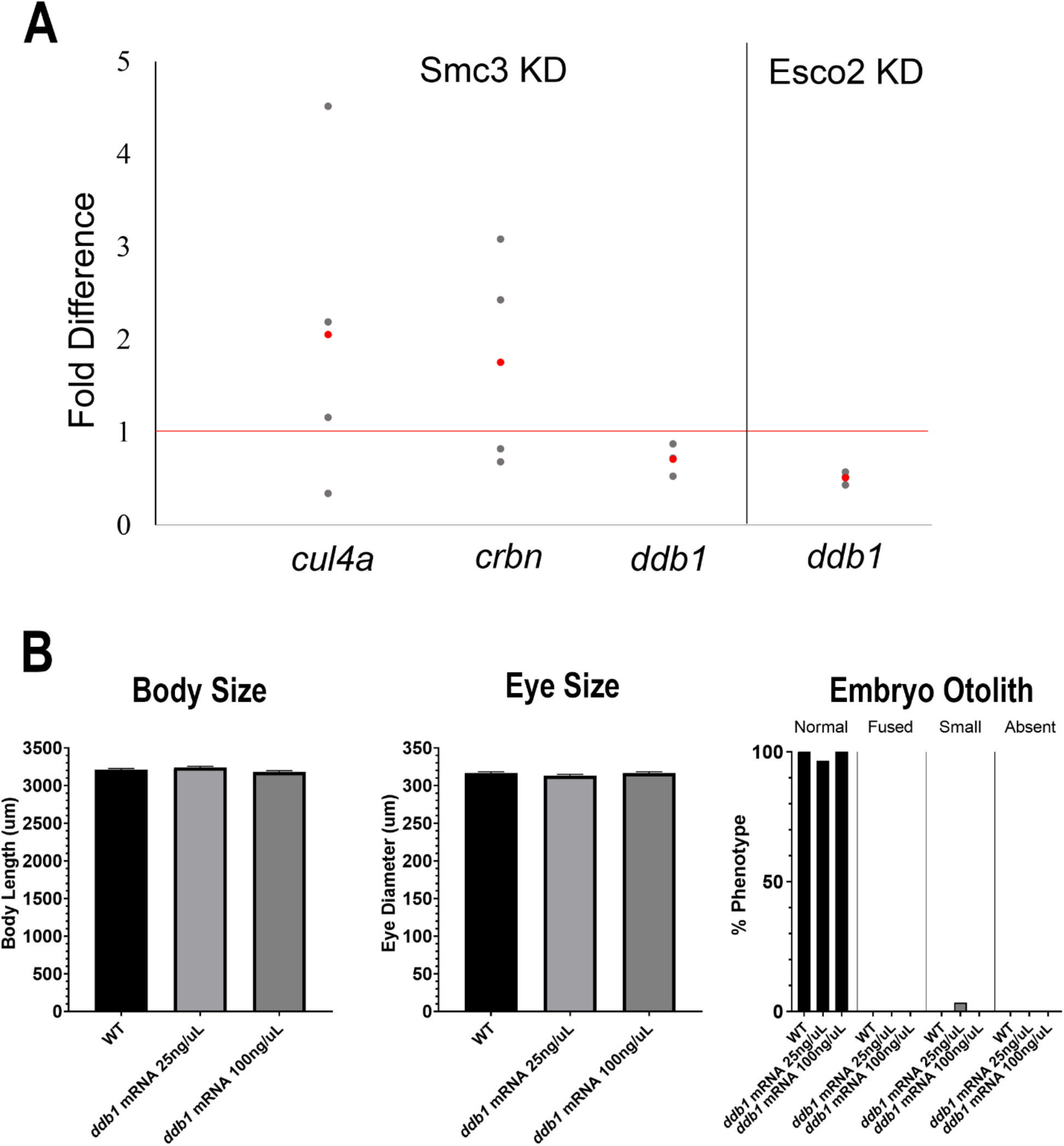
*ddb1* is downregulated in Smc3 KD and Esco2 KD embryos. (A) Gene expression levels were measured by qRT-PCR analysis in Smc3 KD and Esco2 KD embryos. Graph shows fold difference in expression levels from each replicate (grey) and the average expression (red) for each gene. Keratin was used as the internal refence gene control for fold difference calculations. A fold difference of 1 is considered no change with respect to SC MO injected embryos (represented by red line). CRL4 component genes (*cul4a, crbn* and *ddb1*) were analyzed in Smc3 KD embryos. Only *ddb1* expression was significantly downregulated in Smc3 KD (un-paired t-test, P<0.05), while *cul4a* and *crbn* were inconsistent and not significant (un-paired t-test, P>0.05). Esco2 KD embryos also exhibited a significant downregulation of *ddb1* (un-paired t-test, P<0.05). (B) Exogenous *ddb1* mRNA was injected to WT embryos at the 1-cell stage. 30-46 replicates were analyzed and at least 3 independent trials were performed. Phenotypes were imaged and measured at 72 hpf. *ddb1* overexpression in WT embryos had no effect in embryo length, eye size, or embryo otolith phenotype (error bars represent s.e.m, one-way ANOVA with Turkey;s multiple comparison, P>0.05).

**Fig. S3.**
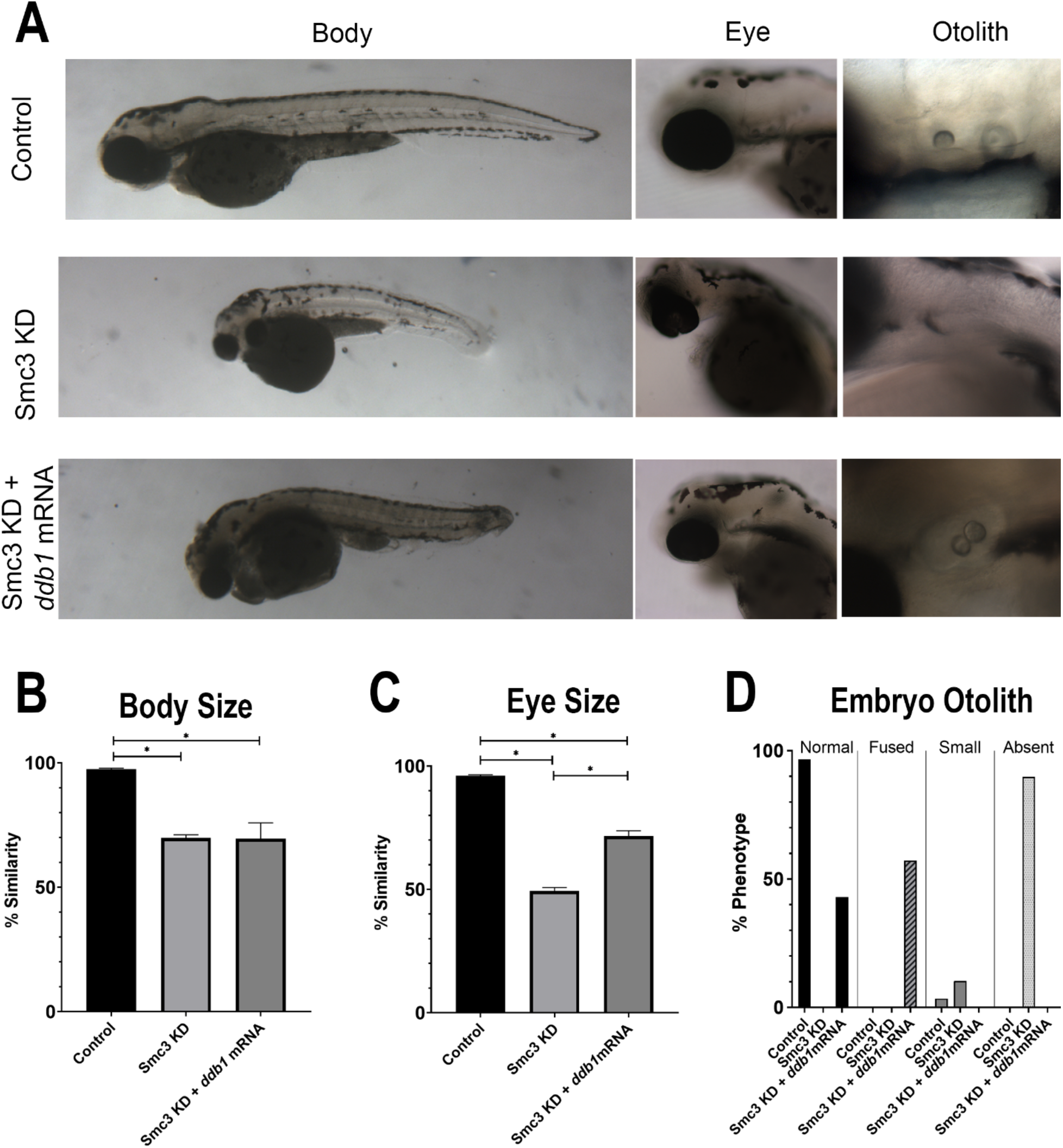
Reduced levels of exogenous *ddb1* overexpression rescues eye and otolith phenotypes in Smc3 KD phenotypes. (A) Representative images of control embryos (WT injected with 25ng/μL concentration of *ddb1* mRNA), Smc3 KD (*smc3*-ATG MO injected) and Smc3 KD +*ddb1* mRNA (*smc3*-ATG MO co-injected with 25ng/μl concentration of *ddb1* mRNA) embryos. For all experiments 40-48 replicates were analyzed and at least 3 independent trials were performed. (B) Quantification of body size from injected embryos were compared to un-injected WT embryos to obtain percent similarity. Bar graph reveals no significant rescue of body length in Smc3 KD +*ddb1* mRNA compared to Smc3 KD alone (error bars represent s.e.m., one-way ANOVA with Turkey’s multiple comparison, P>0.05). (C) Quantification of eye size from injected embryos were compared to un-injected WT embryos to obtain percent similarity. Bar graph reveals a significant rescue of eye size in Smc3 KD +*ddb1* mRNA compared to Smc3 KD alone (error bars represent s.e.m., one-way ANOVA with Turkey’s multiple comparison, *P<0.05). (D) Graph shows percent of normal, fused, small, or absent otolith phenotypes with MO treatments. Data reveals 0% of Smc3 KD embryos exhibited normal otoliths, while 43% of Smc3 KD +*ddb1* mRNA embryo otoliths were rescued to normal levels. A decrease in absent otolith phenotypes was observed with *ddb1* mRNA co-injections compared to KD alone.

**Fig. S4.**
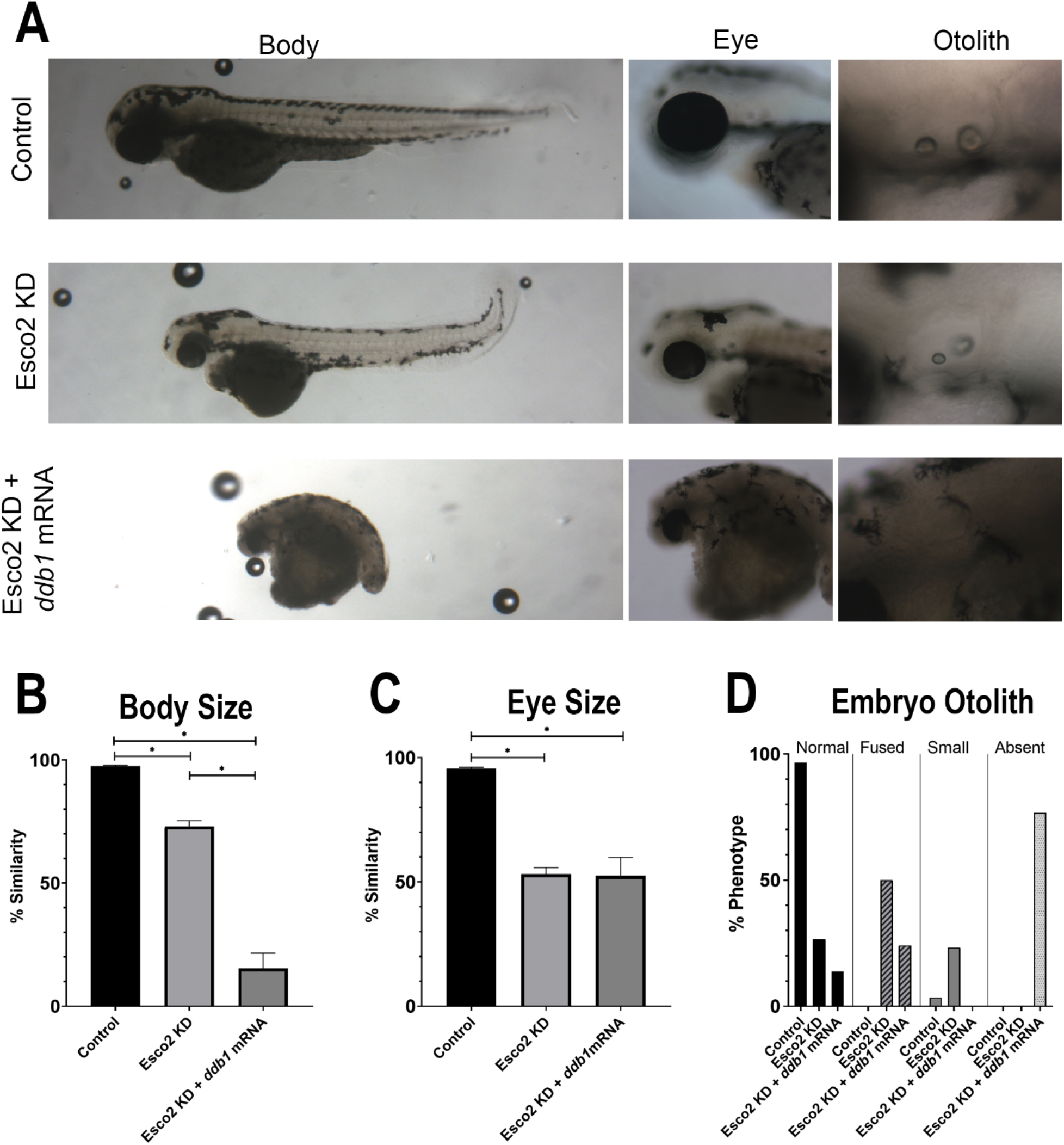
Reduced levels of exogenous *ddb1* overexpression exacerbates body and otolith phenotypes in Esco2 KD phenotypes. (A) Representative images of control embryos (WT injected with 25ng/μL concentration of *ddb1* mRNA), Esco2 KD (*esco2*-ATG MO injected) and Esco2 KD +*ddb1* mRNA (*esco2*-ATG MO co-injected with 25ng/μl concentration of *ddb1* mRNA) embryos. For all experiments 27-30 replicates were analyzed and at least 3 independent trials were performed. (B) Quantification of body size from injected embryos were compared to un-injected WT embryos to obtain percent similarity. Bar graph reveals significant reduction of body length in Esco2 KD +*ddb1* mRNA compared to Esco2 KD alone (error bars represent s.e.m., one-way ANOVA with Turkey’s multiple comparison, *P<0.05). (C) Quantification of eye size from injected embryos were compared to un-injected WT embryos to obtain percent similarity. Bar graph reveals no significant change in eye size in Esco2 KD +*ddb1* mRNA compared to Esco2 KD alone (error bars represent s.e.m., one-way ANOVA with Turkey’s multiple comparison, P>0.05). (D) Graph shows percent of normal, fused, small, or absent otolith phenotypes with MO treatments. Data reveals 27% of Esco2 KD embryos displayed normal otoliths, while only 14% of Esco2 KD +*ddb1* mRNA embryo otoliths exhibited a normal phenotype. An increase in absent otolith phenotypes was observed with *ddb1* mRNA coinjections compared to KD alone.

